# Spatial Regression of Morphology-Protein Coupling in Tumour Proteomics

**DOI:** 10.64898/2026.01.14.699547

**Authors:** Alejandro Leyva, M. Khalid Khan Niazi

## Abstract

Spatial proteomics has enabled high-resolution characterization of protein organization within tumor microenvironments, yet most computational approaches implicitly assume spatial homogeneity and focus on clustering rather than diffusion constraints imposed by tissue morphology. Here, we model morphology–protein coupling in triplenegative breast cancer using geographically weighted regression (GWR) applied to 41 publicly available Multiplexed Ion Beam Imaging (MIBI) samples comprising 36 protein markers. Single-cell morphometric features were extracted from MIBI spots and combined with spatial adjacency graphs to model location-specific protein dispersion. Compared with ordinary least squares and ridge regression baselines, GWR consistently demonstrated superior performance across regression metrics, explaining substantially greater spatial variance in protein intensity (+.4 R^2^ improvements across markers) while reducing mean absolute and squared errors. Information-theoretic analysis showed lower (Aikake Information Criterion Corrected) AICc values for GWR across the majority of markers, indicating improved model fit. Spatial autocorrelation diagnostics further confirmed that GWR residuals exhibited near-random structure, with significant reductions in Moran’s I and Geary’s C relative to global models, demonstrating effective capture of local heterogeneity. Eight proteins with significant spatial autocorrelation, including B7-H3 and -catenin, showed pronounced morphology-dependent dispersion patterns that were not recoverable using global regression. These results demonstrate that explicitly modeling spatial heterogeneity yields more accurate and interpretable representations of protein organization and supports a diffusion-barrier view of pathoproteomics beyond agglomeration alone.

## 1 Introduction

Spatial proteomics examines the visualization and clustering of proteins across histopathological images to understand pathological mechanisms [1]. It provides insight into barriers to drug diffusion, mechanisms behind metastasis and lymphocyte infiltration, and downstream genomic processes [1]. Current models address how spatial clustering of proteins differs between malignant and benign tumors, and how aggregation and content vary across diseased regions [2]. Digital pathologists and bioinformaticians have developed models to predict protein location and concentration within spatial regions, though these approaches are limited by data availability and tend to focus on clustering using k-means neural networks, graphical neural networks, and graphical convolutional networks [2]. A major challenge in spatial proteomics is the assumption of spatial homogeneity across histopathology sections, which overlooks diffusion barriers and paracrine signaling within tumor regions and tissues such as adipose tissue [3]. Protein diffusion is influenced by paracrine, autocrine, and hormonal signaling, but these processes are constrained by diffusivity and membranous barriers [3]. The concentrations and movements of biomarkers relate to histopathological morphology and depend on inherent protein characteristics, including size and hydrophobicity. Modeling spatial heterogeneity therefore provides insight into disease behavior and how tumors proliferate or evade lymphocytic interactions.

Multiplexed Ion Beam Imaging (MIBI) is a spectrometry-based method for evaluating protein concentration at 260 nm resolution across 30 to 40 proteins using metal-bound antibodies [4]. Surgical resections are taken from patients, antibodies are validated for specificity, and then bound to metal ions [4]. After ion-beam exposure, mass spectrometry integrals are used to reconstruct spatial images of protein concentration [4].

In immuno-oncology, spatial proteomics helps characterize immune responses and tumor interactions, supports prognosis, and reveals active signaling pathways within the cellular environment [1]. Spatial arrangement of tumor and immune cells is correlated with prognosis, and identifying proteins and signaling pathways assists in recognizing mutant variants and determining candidates for chemotherapy, immunotherapy, and other treatments [1]. Within the tumor microenvironment, paracrine signaling can indicate T-cell exhaustion and the persistence of immune responses [5]. Spatial proteomics can also identify cytokine signaling and agglomeration within cellular regions, and poor drug perfusion can be explained by protein clustering that impairs uptake [5]. Modeling spatial heterogeneity helps determine the likelihood of proteins aggregating in specific morphological regions, which informs pharmacokinetics [6]. While spatial transcriptomics reveals pathway activation, it cannot show the behavior of the signaling cascade itself. Understanding molecular percolation provides insight into the severity of fibrotic diseases.

Developing models to predict spatial agglomeration of proteins can provide mechanistic understanding of signaling behaviors in tumor-adjacent regions and infer protein concentrations from segmented regions for diagnosis and treatment planning. Spatially informed proteomics models using globally weighted regressions can account for lower diffusivity of proteins across certain tissue regions.

## 2 Materials and Methods

41 patient slides from TNBC were imaged using MIBI across 36 different proteins for triplenegative breast cancer, and all data are publicly available [1]. MIBI images were preprocessed using image rescaling to ensure all protein markers were captured using contrast and brightness adjustments. Each individual sample was coupled with 36 corresponding images labeled with the specific protein tracked by the ion beam. Centroid graphs using k-nearest neighbors and Delaunay triangulation were constructed to analyze protein clustering across regions. An ordinary least squares model was used as a baseline for comparison.

All MIBI marker images were normalized using robust percentile rescaling, where pixel intensities were clipped to the 1st and 99th percentiles and scaled to the 0–1 range. Cell segmentation masks were imported from labeled TIFF files [1], and centroid coordinates were extracted for each cell. For every marker, per-cell intensity features were computed using thresholding based on Li’s method with an Otsu fallback [7, 8]. The features chosen include the fraction of positive pixels, integrated density, mean intensity of the top five percent of pixels, maximum intensity, Gini coefficient of intensity distribution, and the number of positive-signal blobs, along with log and square-root transforms. Morphological features (area, perimeter, eccentricity, solidity, extent, equivalent diameter) were z-scored and reduced by PCA to obtain a single MorphScore capturing dominant shape variation. Centroid coordinates were used to construct a spatial adjacency graph via Delaunay triangulation with a k-nearest neighbor fallback to ensure connectivity. Edges were constructed based on spatial proximity and were minimally encoded. Moran’s I and Geary’s C were computed for each marker’s log-transformed intensity using permutation-based significance testing [9, 10], but spatial weights were constructed separately by the geographically weighted regression analyses [10].

Only the top eight markers (B7H3, -catenin, CD11b, CD11c, CD138, Au, TSPAN8, and SPINT2) were backpropagated for globally weighted regression, and only markers with a Moran’s I significance above 0.05 were used in the model. Proteins were included only if they demonstrated residual spatial autocorrelation under the k-nearest neighbor graph. Cellular metrics including eccentricity, solidity, identity, extent, and equivalent diameter were recorded where proteins were detected to account for spatial features of cells. Delaunay graphs were constructed over cell centroids at protein-positive locations and used exclusively for spatial diagnostics. Protein concentration was modeled as a function of cell-level morphometric features, with spatial heterogeneity captured through geographically weighted regression, allowing regression coefficients to vary continuously across tissue coordinates..

Finally, because MIBI images form a set of Voronoi-like tessellations, Delaunay triangulation provides an appropriate spatial arrangement of centroids for downstream modeling. **Notation**. For each observation *i* = 1, 2, …, *n*:

- *y*_*i*_ – dependent (response) feature at subcellular location *i*;
- *x*_*ik*_ – value of the *k*^th^ explanatory variable at location *i*;
- (*u*_*i*_, *v*_*i*_) – spatial coordinates (e.g., centroid or pixel coordinates) of observation *i*;
- *β*_0_(*u*_*i*_, *v*_*i*_) – spatially varying intercept term;
- *β*_*k*_(*u*_*i*_, *v*_*i*_) – location-specific regression coefficient for the *k*^th^ predictor;
- *ε*_*i*_ – random error term, assumed to follow *ε*_*i*_ ∼ 𝒩 (0, *σ*^2^);
- *X* – *n* × *p* design matrix of predictors;
- **y** – *n* × 1 response vector;
- *W* (*u*_*i*_, *v*_*i*_) – diagonal spatial weighting matrix for location (*u*_*i*_, *v*_*i*_);
- *w*_*ij*_ – spatial weight quantifying the influence of observation *j* on location *i*;
- *d*_*ij*_ – Euclidean distance between locations *i* and *j*;
- *b* – bandwidth parameter controlling the spatial kernel size.

The regression task uses coefficients that change from one spatial location to another, so the influence of each morphometric feature is allowed to vary across the tissue. The unexplained portion of the model is captured as residual variation, and the intercept at any location is simply the locally estimated baseline value produced by the regression at that position.

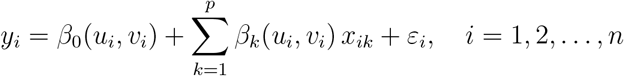

Ordinary least squares estimates the location-specific coefficients by applying the spatial weights to the morphometric features and fitting them to the observed outcomes.

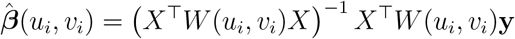

The weight matrix is formed as a diagonal matrix in which each entry represents the weight assigned to a specific observation relative to the spatial location being evaluated.

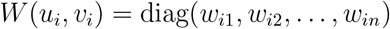

Each location’s weights are generated by applying an exponential decay to the Euclidean distance between points, controlled by the bandwidth parameter. When a point falls within this bandwidth, its weight reflects how close it is to the target location, and points beyond this distance are assigned a weight of zero so that only nearby observations influence the local regression.

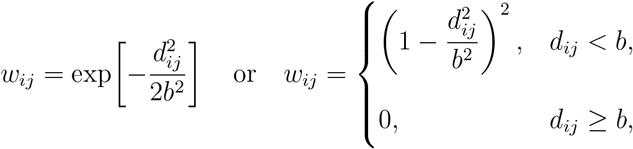

The output is obtained by combining the morphometric features with the spatially weighted regression fit, along with the residual error. The corresponding expression for the spatial regression estimate follows directly from the ordinary least squares formulation.

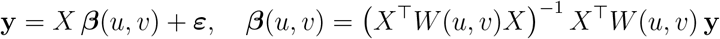

## 3 Results

The results for the regression tasks demonstrated superior performance of geographically weighted regression across all metrics against the baselines. Ridge and Unaltered OLS had a signifcant but erroneous predictive capacity, only explaining around 4% of variance for given marker’s dispersion. Modelling spatial heterogeneity around each instance provides inference on the likelihood of dispersion within a cell.

Figure 2 shows the distribution of results for the R squared for each biomarker, demonstrating that across the 40 markers, Geographically weighted regression explains higher spatial variance for predictive resgressions. Aikake information criterion was measured between both OLS and GWR, though Delta AIC proves that OLS had a much higher OLS on average across multiple markers predicted at each point. The distribution of AICcs when compared against OLS shows that the majority of values center toward zero, likely due to low dispersion of certain markers. Ultimately, Geographically weighted regression captures spatial relationships in protein dispersion that aren’t conclusive by agglomeration alone.

**Figure 1.**
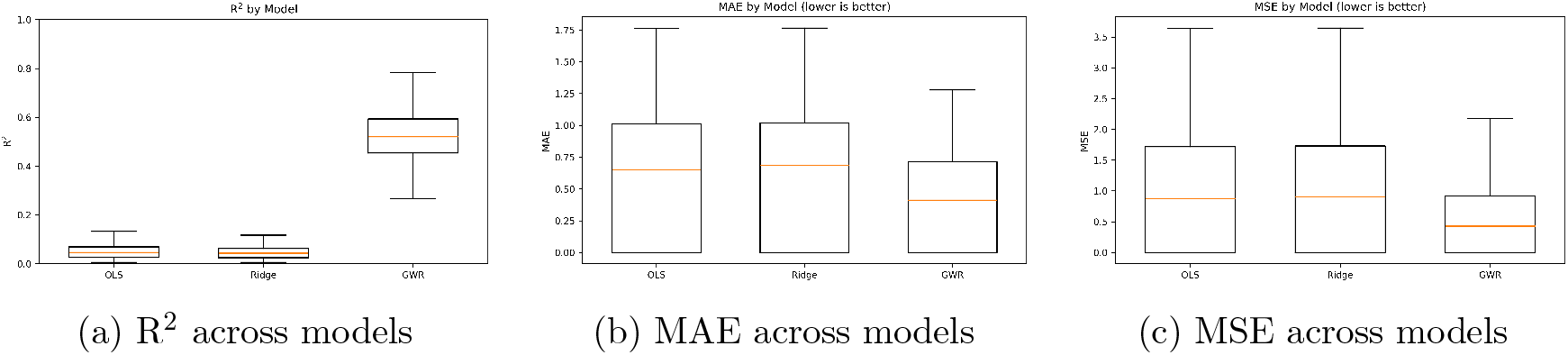
Comparison of global (OLS, Ridge) versus local (GWR) regression performance. GWR shows higher *R*^2^ and lower MAE/MSE, indicating improved fit to spatially heterogeneous data.

**Figure 2.**
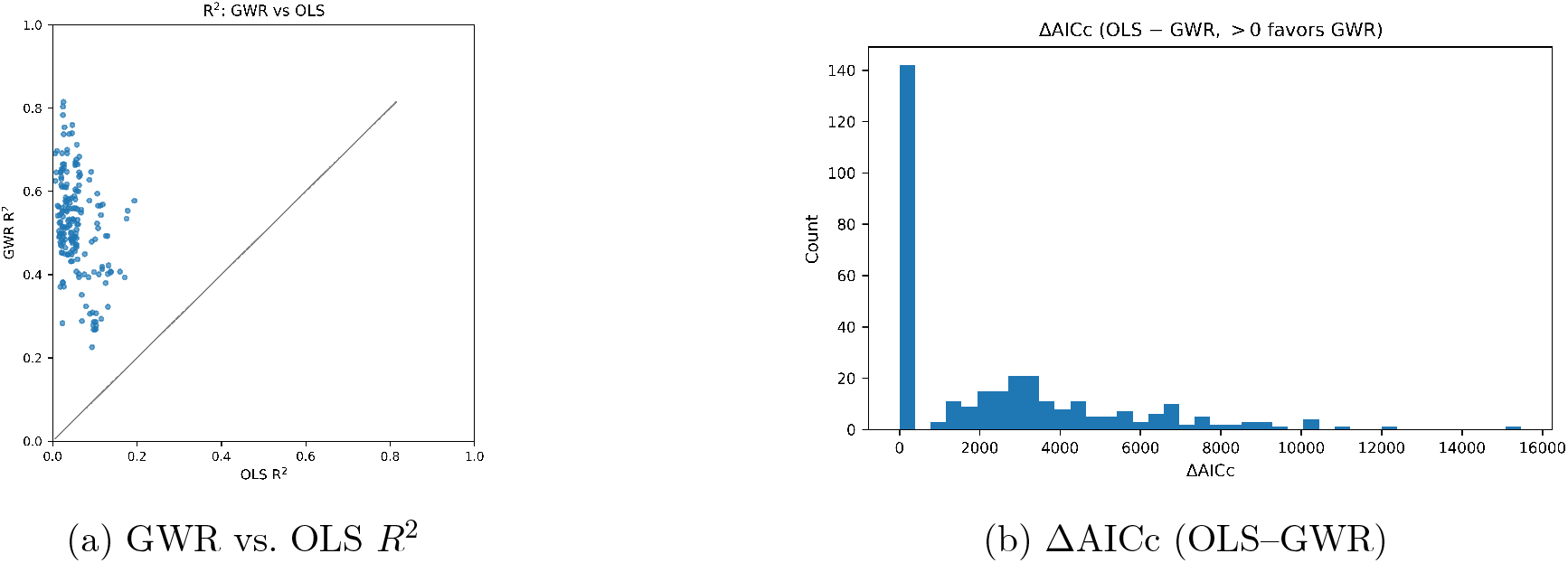
Information-theoretic comparison of model fit. Positive ΔAICc and diagonal shift toward lower AICc confirm superior fit of geographically weighted regression.

The residual Moran’s index shown in Figure 3a. shows that OLS is superior at capturing global clustering trends. However, GWR clearly captured Residual Geary’s index in (c) with avidly higher performance then OLS, proving that the GWR model can capture local heterogeneity in tissue. Figure b and d demonstrated that the distribution of values for gwr were much higher and lower respectively.

**Figure 3.**
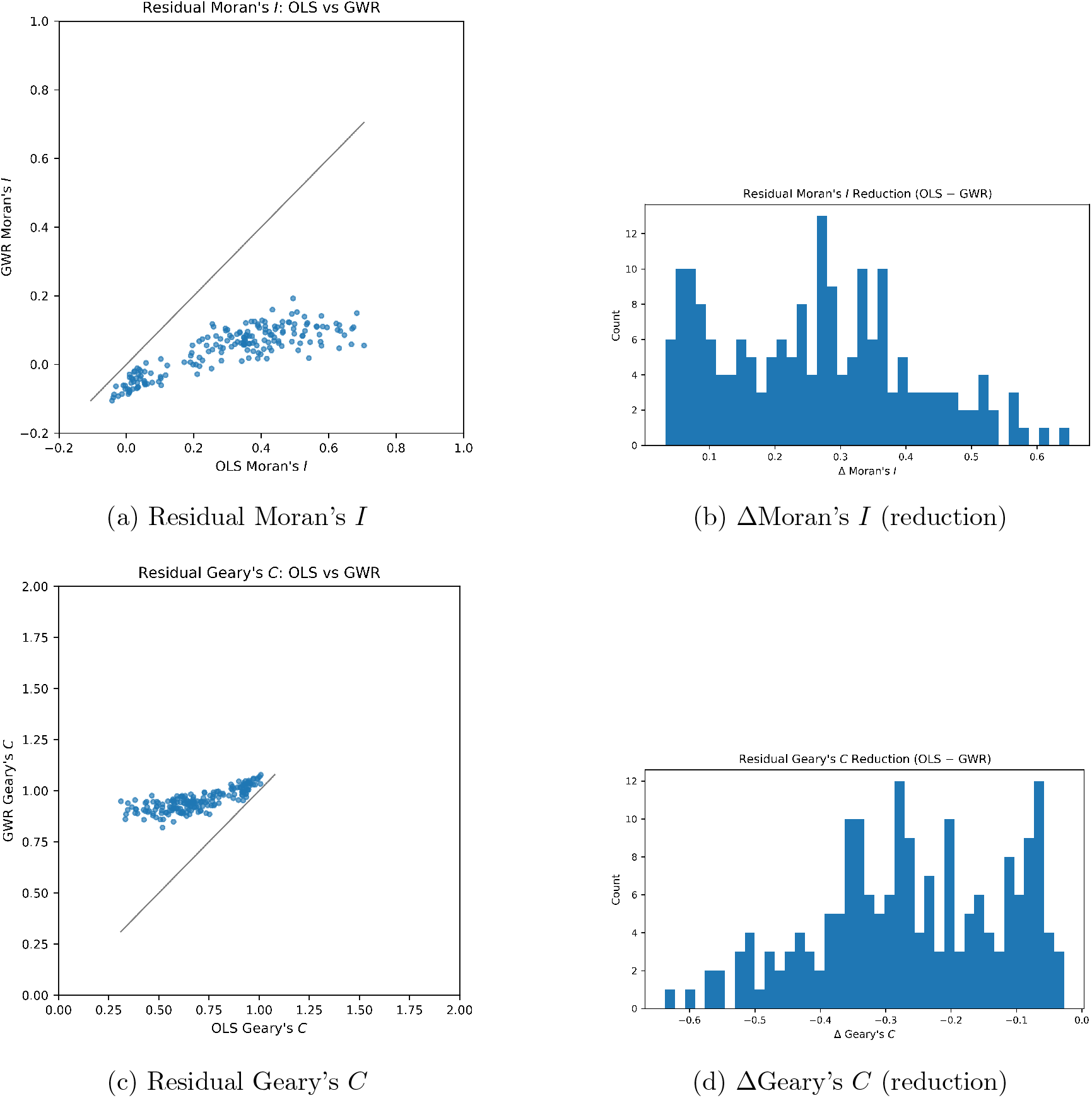
Spatial autocorrelation diagnostics before and after local modeling. Residuals from GWR exhibit near-random spatial structure, confirming that local terms capture spatial dependence.

**Figure 4.**
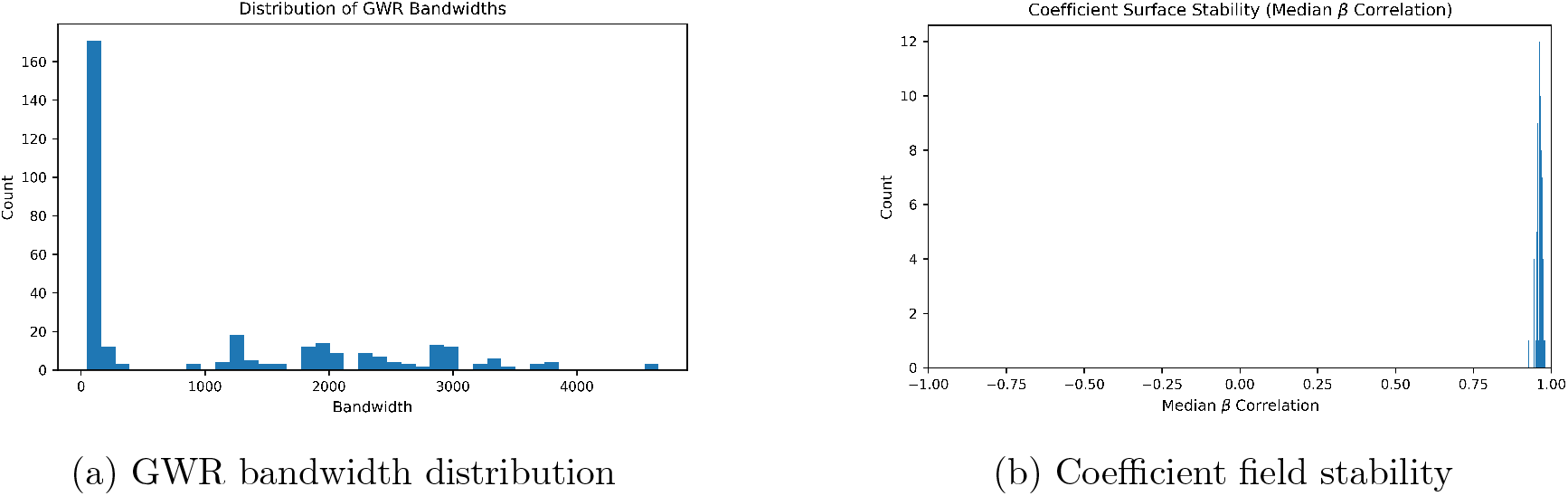
Local bandwidth variability reflects multiscale spatial processes, while high median *β*-field correlation indicates stable coefficient surfaces across the tissue.

The distribution of GWR bandwidths had a high count of markers with a spatial band-width slightly above zero because a variety of signficant markers did not display any significant dispersion. Tissue coordinates were in pixel units; with AICc-based selection this can favor very small kernels. In this run, many markers converged at the minimum feasible bandwidth, indicating a boundary solution of the selector rather than a biological hard scale. The correlationn heat map shows a high correlation with cellular features and biomarkers, though certain biomarkers are not as prevalent and thus lowly correlated.

The distribution of values in figure 6 for the gold markers and beta catenin biomarkers demonstrates clustering around regions, and GWR is capable of capturing spatial heterogeneity around these regions. This can be used to understand the clusterings of cell regions using spatial heterogeneity coupled to traditional clustering algorithms and deep learning modules. In addition to understanding pararcine signaling between molecules, the perfusion of molecules and markers can be understood through the lens of diffusive barriers rather than agglomeration.

**Figure 5.**
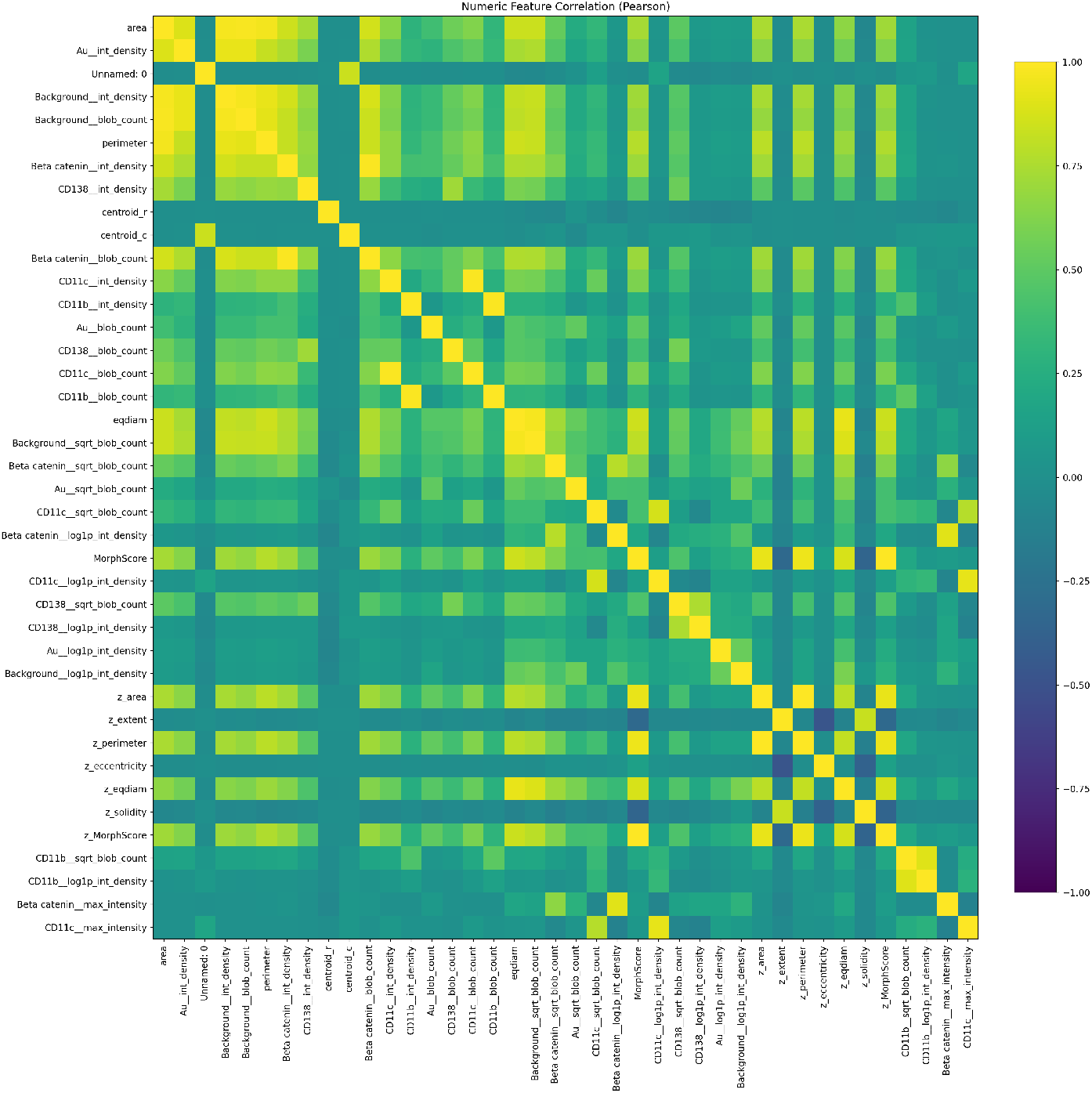
Correlation matrix of single-cell morphological and marker-intensity features, revealing structured co-variation within the tumor microenvironment.

**Figure 6.**
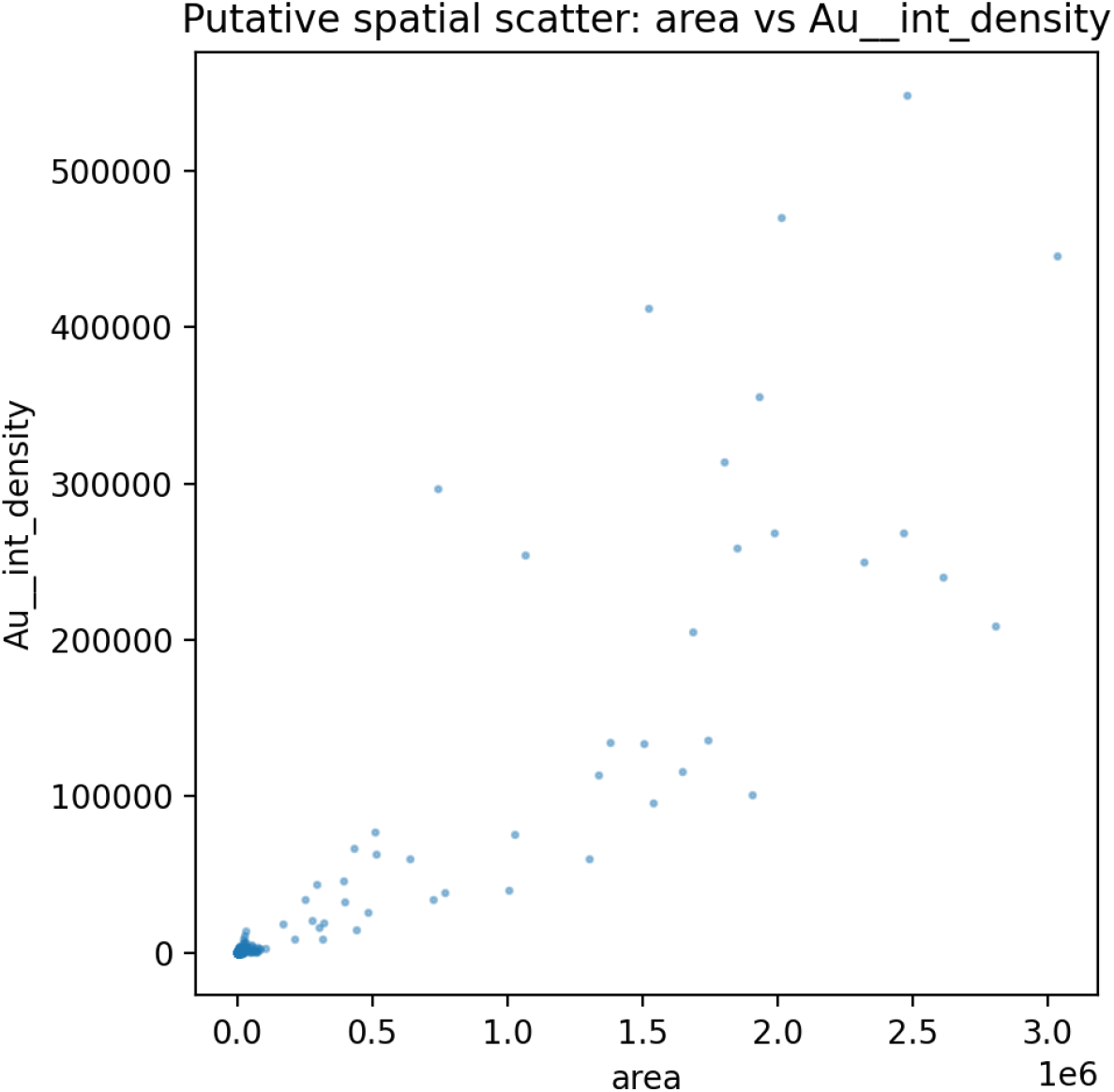
Spatial scatter of cell area versus Au intensity as a representative spatial feature map. Localized clusters illustrate heterogeneous microenvironments captured by GWR.

**Figure 7.**
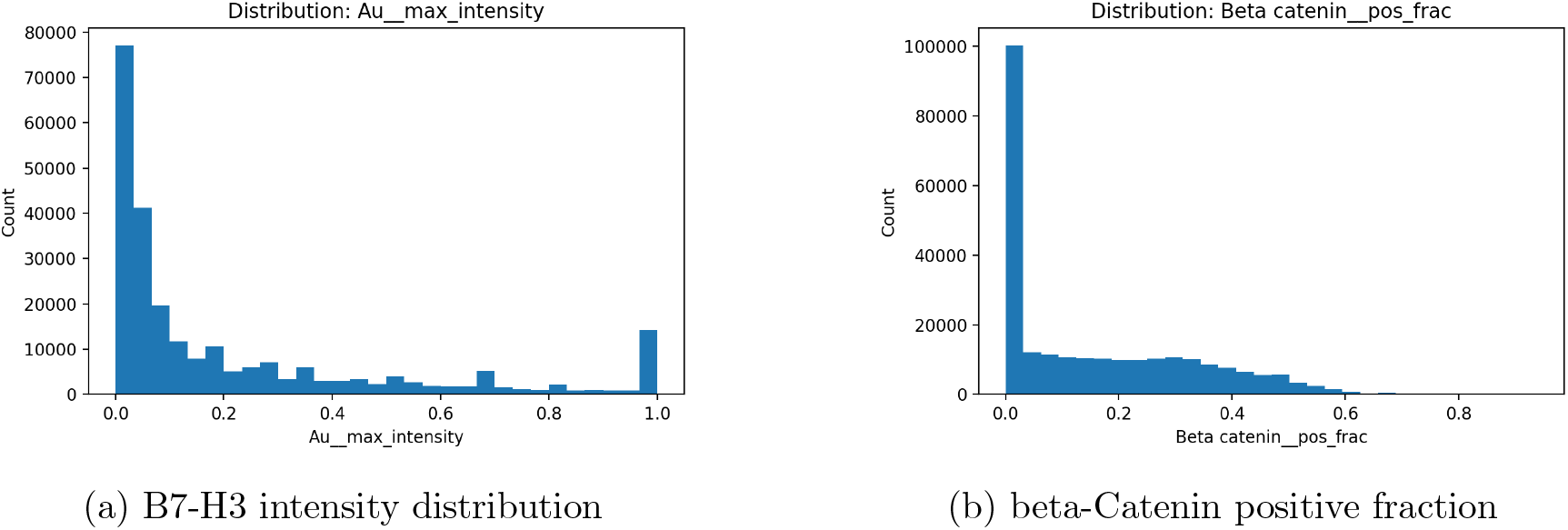
Single-cell distributions of B7-H3 and beta-Catenin reveal strong intercellular heterogeneity across the tissue section.

The Gold biomarker density that was coupled to antibodies was measured to determine the spatial heterogeneity and perfusion of these markers across a tissue surface. Understanding spatial coupling and location perfusion barriers can be used to understand the perfusion of nanomedicine based therapies and small molecule drugs.

The distribution of molecules across these instances provides the likelihood of protein signaling and clustering with regards to spatial barriers and morphological consistencies. GWR has proven to be useful for spatial proteomics and general pathoinformatics.

## 4 Discussion

Spatial proteomics provides a means to understand the clustering of proteins around immunocellular and endothelial–epithelial barriers[1,2]. Globally weighted regression offers a proven approach for modeling spatial barriers and heterogeneity across cellular and tissue structures, enabling applications in precision therapies, pharmacology, and drug delivery[5]. With the emergence of optical nanoparticles and nanoscale delivery systems, modeling the spatial structure of tumor sites becomes valuable for predicting perfusion efficiency and treatment penetration.

Traditional deep learning methods such as graphical neural networks primarily characterize cluster distributions or local neighborhoods, but they do not explicitly model spatial barriers that influence protein organization within metastatic tumors[8]. By incorporating spatial heterogeneity through geographically weighted regression, the mechanisms underlying protein dispersion and localization can be more accurately inferred, providing deeper mechanistic insight into tumor microenvironment behavior.

## 5 Limitations

The algorithm is limited by the dispersion of proteins across celullar channels, and if less proteins are dispersed or agglomerated, then the model will fail to predict the likelihood of clustering around certain instances. It is crucial to note that geographically weighted regression accounts for linear relationships. Non-linear regression such Geographically neural network weighted regressions were not tested due to the heterogeneity of protein tasks and the image processing required to integrate. This study serves as a proof-of-concept that geographically weighted regressions can be used to understand the spatial dispersion of proteins with respect to morphology.

## 6 Conclusions

Geographically weighted regression outperforms traditional regression tasks for predicting spatial autocorrelation of protein dispersion across MIBI images. this implies that spatial heterogeneity is a valuable variable to accomodate when when accounting for measuring protein agglomeration. For future work, this study can be expanded to incorporate graphically weighted geogrpahical neural networks to account for the nonlinear relationships accounting for protein clustering and spatial heterogeneity. This method can also be translated into other methods of spatial proteomics images, such as MALDI TOF. Spatial regression provides a new and interpretable method of predicting spatial correlation of proteins based on morphology.

## 7 Ethics Statement

All datasets are public and anonymized.

## 8 Author Contributions

All work was performed by the first author, with supervision by the second.

## 11 Data Availability

The data is from citation 1, and is a publically available MIBI protein dataset for triple negative breast cancer. All code is available upon request. We would like to acknowledge the superb work of Dr, Keren’s group at the Weizzman institute. In addition, the code is posted here: https://github.com/Alejandro21236/SpatialProteomicsGWR

